# Dynamics of Expression Variability Underpin Retention of Small-Scale vs. Whole-Genome Duplicates

**DOI:** 10.1101/2024.11.06.622370

**Authors:** Haoran Cai, David L. Des Marais

## Abstract

Genome analyses reveal that gene duplication in eukaryotes is pervasive, providing a primary source for the emergence of new genes. However, the mechanisms dictating early duplicate retention and the emergence of functional biases—such as the enrichment of tandem duplicates in environmental responses—remain unclear. Here, to better understand the mechanisms and factors determining gene retention, we study a frequently overlooked molecular feature—expression variability—as measured by within-line expression variation, termed variability. We demonstrate that, on average, genes with duplicates exhibit higher expression variability than singletons. Furthermore, small-scale duplications (SSDs) and whole-genome duplications (WGDs) display contrasting functional outcomes and time-dependent profiles in expression variability. These findings suggest a potential overarching mechanism that facilitates gene expression divergence, functional gains of environmental responses, and duplicate retention following SSDs.

## Introduction

Gene duplication is recognized as the primary source of new genes and gene functions and has consequently been credited with great evolutionary importance (Ohno 1970, Des Marais and Rausher 2008). Typically, gene duplication generates two gene copies; this theoretically allows one or both copies to evolve under reduced selective constraint and, on some occasions, to acquire novel gene functions that contribute to adaptation.

Duplicate gene copies can be generated through one of two broad mechanisms, namely small-scale or large-scale duplication events. Among the most dramatic forms of large-scale duplication is whole-genome duplication (WGD), which leads to a sudden expansion in genome size and, significantly, preserves the chromosomal context of most genes (Kuzmin et al. 2022). WGDs have likely occurred multiple times throughout 200 million years of angiosperm evolution (Vision et al. 2000). Conversely, small-scale gene duplications (SSD), including proximal duplication, tandem duplication, and transposed duplication, result in duplication of a specific genomic region, often comprising a single gene (Wang et al. 2012*b*, Kuzmin et al. 2022, Zhang 2003). Plant genomes, on average, harbor duplicate copies for approximately 65% of annotated genes. Most duplicates originate from WGD events and, accordingly, paleopolyploidization has been cited as an important source of evolutionary novelty for land plants (Panchy et al. 2016, Tiley et al. 2016). While WGD events lead to a sudden and massive expansion of the genome, they are often followed by extensive gene loss over evolutionary time (Lynch and Conery 2000). Nevertheless, WGD-derived duplicates tend to be retained at a higher rate compared to SSD-derived duplicates (Panchy et al. 2016).

Considerable evidence suggests that duplicates originating from WGD and SSD events differ in their evolutionary rate, essentiality, and function (Qiao et al. 2019, Freeling 2009). For example, SSD duplicates are more prone to neo-functionalization whereas WGD duplicates tend to partition ancestral functions (Fares et al. 2013). This difference is apparent in the functional similarity observed among WGD paralog pairs compared to SSD paralogs (Hakes et al. 2007). In addition, models of gene and genome duplication suggest that the rate of gene retention following duplication is notably different for WGD and SSD events, with biases towards specific functional classes (Maere et al. 2005, Freeling 2009). Particularly, genes originating from SSDs often play roles in environmental and defense responses, while genes retained from WGDs are associated with intracellular regulation and include transcription factors, ribosomal proteins, and other core cellular processes. (Hanada et al. 2008, Rizzon et al. 2006, Wang et al. 2012b, Qiao et al. 2019, Seoighe and Gehring 2004). However, the underlying mechanisms driving such functional biases are not well understood. As such, we remain uncertain whether genes retained following the different duplication sources are subjected to different evolutionary constraints, or whether the two classes of genes possess equivalent potential to generate novel function. The gene balance theory offers one plausible explanation for why WGD events predominantly retain transcription factors (Freeling 2009). It has been hypothesized that the more connected the gene product, the more likely the phenotype will change if dosage imbalance occurs. Therefore, the gene balance theory posits that if a tandem duplicate is highly connected, it is more prone to cause dosage imbalance and thus is very likely to be removed. Conversely, after WGD, loss of a highly connected duplicate would be more likely to cause dosage imbalance and deleterious effects and thus is more likely to be retained (Wang et al. 2012b, Freeling 2009).

Theoretical and empirical results show that most duplicated genes return to single copies after duplication because a functionally redundant duplicate will accumulate deleterious mutations and evolve into a pseudogene (Wagner 1998, Tautz 1992, Lynch and Conery 2000). Therefore, the earliest stage following duplication is key to understanding the role of gene and genome duplication in evolution (Innan and Kondrashov 2010). Classic models of duplicate retention include neofunctionalization (Kuzmin et al. 2022, Ohno 1970), subfunctionalization (e.g., the duplication–degeneration–complementation model, DDC) (Van Hoof 2005, Marshall et al. 2013), and escape from adaptive conflict (a special case of subfunctionalization)(Des Marais and Rausher 2008, Kuzmin et al. 2022).

Changes in gene expression can play an important role in the preservation of duplicated genes (Francino 2005, Ganko et al. 2007, Duarte et al. 2006, Doebley and Lukens 1998, Casneuf et al. 2006, Duarte et al. 2006). One observation is that gene expression exhibits greater divergence for pairs originating from SSD as compared to WGD (Casneuf et al. 2006), which may arise from a higher likelihood of altering regulatory features during the process of SSD (e.g., if the duplication event does not faithfully replicate *cis*-regulatory elements) (Rogers et al. 2017). Although seemingly contradictory to such observations, WGD pairs, in fact, have a higher average number of *cis*-element differences between paralogs than do SSDs (Arsovski et al. 2015). Thus, a question remains as to whether changes in transcription contribute to the probability of gene retention and the subsequent functional biases observed in retained paralogs. Another relevant and intriguing question is whether variants affecting gene expression dosage experience the same degree of relaxed selection as do variants that affect protein coding sequences; it is generally accepted that functional redundancy of a protein-coding gene leads to relaxed selection constraints (Ohno 1970, Kondrashov et al. 2002). If this is also the case with transcriptional expression, it would naturally increase the ‘search space’ of a new copy, and potentially facilitate gene retention via neofunctionalization.

Here, we study expression variability among gene duplicate pairs to test the hypothesis that biases in the functions of gene copies retained can be explained by changes in regulatory architecture immediately after duplication. Expression variability is defined as the variation of gene expression among genetically identical individuals in a controlled environment (Cortijo et al. 2019, Cortijo and Locke 2020). In multicellular organisms, such as plants, genetically identical or clonal individuals are used to assess variability, also referred to as inter-individual variability (Cortijo et al. 2019) or intra-genotypic variability (Metcalf and Ayroles 2020). In the context of gene duplication, a duplicate may change its genomic context, via transposition or incomplete duplication in which regulatory elements are reshuffied (Arsovski et al. 2015), thereby changing expression variability of the new copy as it becomes fixed through neofunctionalizaion. Here, we first show that duplicates arising from both SSDs and WGDs tend to exhibit elevated expression variability. Our subsequent analyses point to distinct mechanisms that could explain the increasing expression variability observed for SSDs and WGDs.

## Materials and Methods

### Normalized expression variability

Gene expression data was obtained from Cortijo et al. 2019. Briefly, RNA-sequencing of 16 individual seedlings of *A. thaliana* (Col-0 accession) were used to calculate by-gene inter-individual expression variability. The square coefficient of variation (*CV* ^2^ = *variance/*(*average*^2^)) is first calculated. To account for the coupling between the mean gene expression level and *CV* ^2^ and to allow for comparisons among genes, we calculated the normalized expression variability (NEV) as log2(*CV* ^2^/trend for the same expression level)(Cortijo et al. 2019).

### Calculation of environmental responsiveness from expression atlas

We used a large gene expression atlas (Roberts and Josephs 2023), encompassing a range of experimental conditions and tissues, to characterize the plastic expression responses of individual genes. To calculate environmental responsiveness, we focused on the ‘Col-0’ ecotype and ‘seedling’ tissue across environmental treatments (e.g., cold exposure, day length, hormonal treatment) and developmental stages in the atlas. Responsiveness was quantified as the sum of log2 ratios between expression in a treatment and a baseline condition:

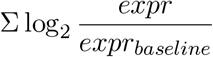

### Tissue specificity

Tissue specificity of individual genes were obtained from Yang and Gaut 2011, which were calculated from Arabidopsis Development Atlas (ADA), and from the expression atlas (Roberts and Josephs 2023). The expression level of a gene was estimated by the average value of all 79 samples in Roberts and Josephs 2023. The tissue specificity was measured with the index *τ* (Yanai et al. 2005):

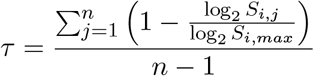

where *n* is the number of tissues, *S*_*i,j*_ is the expression of gene *i* in tissue *j*, and *S*_*i,max*_ is the highest expression of gene *i* across *n* tissues. The index *τ* ranges from 0 to 1, with a higher value indicating a higher specificity. If a gene is expressed in only one tissue, *τ* approaches 1.

### Paralog pairs

The designations of genes as duplicates or singletons were acquired from Yang and Gaut 2011 according to the assignments of Blanc et al. 2003. Paralogs and their corresponding duplication modes were obtained from Coate et al. 2020 and Wang et al. 2013. *k*_*s*_ were similarly obtained from Coate et al. 2020, where alignments of protein sequences of duplicate genes were produced using ClustalW (Thompson et al. 1994).

## Results

Following Cortijo et al. 2019, we first obtained normalized expression variability for each gene at each time point during a day. Gene expression data were acquired from 16 seedlings of the Col-0 wild type of *Arabidopsis thaliana*. The goal of normalization is to account for the effect of coupling between mean expression and variation around the mean, with the aim to compare expression variability across genes. Hereafter, we use median normalized expression variability among all time points to quantify a focal gene’s expression variability, termed normalized expression variability (NEV).

### Elevated gene expression evolvability for duplicates

We first categorized genes into three groups using their duplication status acquired from Yang and Gaut 2011: singletons (i.e., no known duplicates in the *A. thaliana* genome), duplicates arising from WGD, and duplicates arising from SSD (Fig. 1A, Fig. S2, and Fig. S3). Despite the observation that the average NEV effect size is small, the distributions of NEV of duplicates of either type are significantly shifted compared to genes without duplicates (Fig. 1A and Fig. S3, adjusted p = 0.0007 with SSDs, 0.047 with WGDs). We detect no significant difference in NEV between SSDs and WGDs (Fig. 1A and Fig. S3, adjusted p = 0.27). We also conducted similar analyses by stratifying the data based on specific features of genes (Fig. S1) and gene ages (Fig. S2). Further analysis reveals that this elevated expression variation is primarily driven by divergence in NEV between paralogs, rather than uniformly high NEV across all duplicates (Fig. 1B).

**Figure 1:**
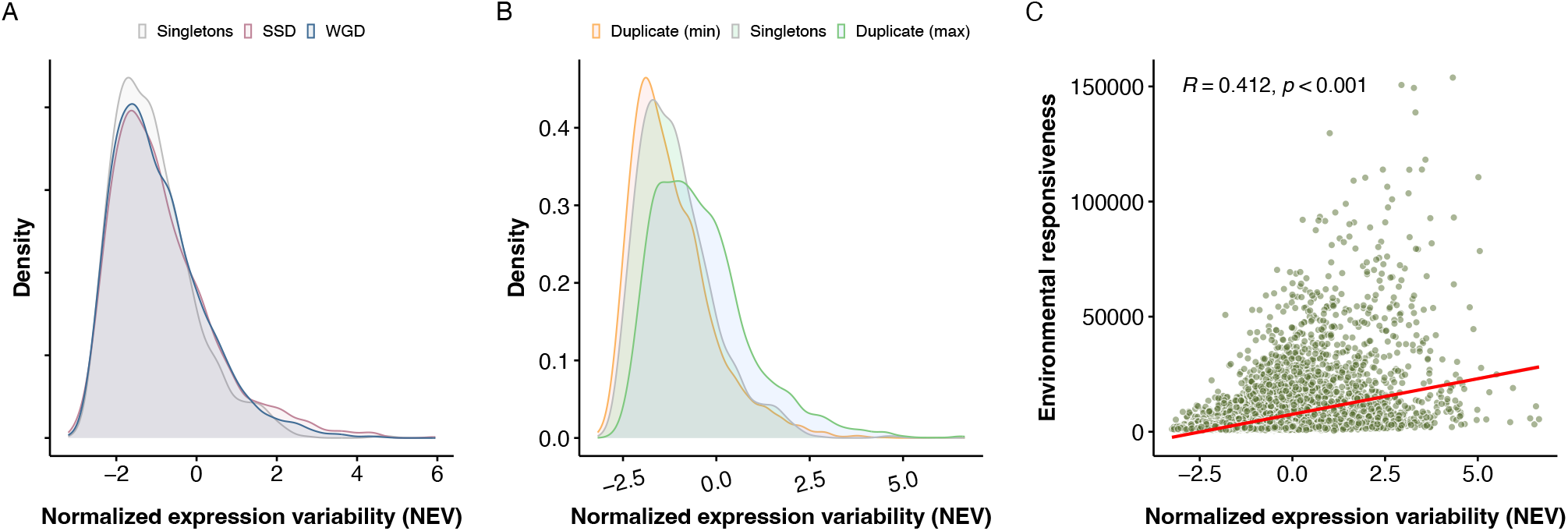
Overall properties of normalized expression variability (NEV) A). Distribution of NEV grouped by singletons (N = 1241), genes arising from SSD (N = 3366), and from WGD (N = 3175). Genes born from whole genome duplications (WGDs) and small-scale duplications (SSDs) exhibit significantly higher NEV than singletons (Wilcoxon two-sided test adjusted p = 0.047 and Cliff’s Delta effect size = 0.047 95% CI [0.010, 0.084] for WGDs vs singletons, Wilcoxon test adjusted p = 0.00071 and Cliff’s Delta effect size = 0.070 95% CI [0.034, 0.106] for SSDs vs singletons, Wilcoxon test ajusted p = 0.27 and Cliff’s Delta effect size = 0.024 95% CI [-0.004,0.052] between SSDs and WGDs). B). Distributions of NEV grouped by genes with lower NEVs within a paralog pair, singletons, and genes with higher NEVs within a paralog pair. N = 1241 for singletons, N = 4344 for each duplicate group (max and min) C). The plasticity of transcriptional responses (responsiveness) summarizes the ability of a gene’s transcript to vary in response to a wide range of environmental perturbations. In our case, we obtained an expression atlas and calculated responsiveness using expression data in seedlings and col-0 ecotype of *Arabidopsis* (Roberts and Josephs 2023). Normalized expression variability (NEV) correlates with plasticity of transcriptional responses. Numbers in the plot are the Pearson correlation coefficient and associated p-value

### Expression variability as an evolvability index for gene expression

We next ask whether expression variability is coupled with plasticity caused by macro-environmental variation, finding that genes with higher NEV tend to show higher sensitivity to environmental perturbations (Fig. 1C). Conversely, genes with lower NEV are more often insensitive to environmental perturbation. While one of the drivers of NEV is micro-environmental variation among individuals, such differences are small, random, and distinguishable from macro-environmental variation that characterizes systematic differences of environmental parameters in experimental settings (Masel and Siegal 2009). This distinction is apparent as indicated in Fig. 1C: higher NEV does not necessarily lead to higher sensitivity to environmental perturbations in an experiment, while a lower NEV constrains the environmental responsiveness of a gene (the apparently inaccessible top left area in the scatter plot of Fig. 1C).

Notably, such noise–plasticity coupling has been observed in single cell organisms, and has been interpreted as a constraint on the evolution of gene expression more broadly (Lehner 2010, Singh 2013, Chapal et al. 2019). We also observe a positive correlation between tissue specificity and normalized expression variability in two independent datasets, which were calculated from Arabidopsis Development Atlas (ADA) (Yang and Gaut 2011), and from the expression atlas (Roberts and Josephs 2023) (Fig. S4).

Prior work on the effects of naturally occurring mutations on gene expression found that genes with greater sensitivity to mutation exhibit higher stochastic noise in gene expression and are also more sensitive to environmental perturbation (Landry et al. 2007). In single-cell organisms, stochastic noise describes the variation within a population of genetically identical cells under a single environment. Together with our study, these findings in yeast mutation accumulation lines imply that expression variability may play a role in the evolution of environmental responses more broadly. We thus hypothesize that elevated expression variability increases the probability of gene retention following duplication, particularly through functional gains of environmental responses.

### Contrasting functions among SSD and WGD paralogs despite similar levels of NEV

We conducted GO enrichment and pairwise semantic similarity analyses on pairs of duplicated genes that show the greatest divergence in NEV between copies (the 5% of genes exhibiting the greatest divergence, comprising 116 SSD pairs and 101 WGD pairs, Fig. 2A and Fig. S7). The semantic similarity analyses indicate that diverged SSD pairs are more likely to exhibit different functions between paralogs than pairs arising from WGDs (Fig. 2B and two additional similarity measures in Fig. S8). Next, we specifically tested whether the “high NEV” paralogs from each pair, as a group, exhibit different functional enrichments than the group of “low NEV” paralogs (Fig. 2C, Supplemental tables 1-6). For SSDs, genes with higher NEV in a SSD paralog pair show distinct functions as compared to the lower group. In particular, paralogs in the SSD high group are enriched for functions in biotic and abiotic environmental responses such as responses to cold, zinc ion, fungi, and herbivores. Previous studies found that genes derived from small-scale duplications (SSD) are more likely to be involved in response to environmental stimuli (Hanada et al. 2008). Here, we provide further evidence that, for SSDs, abiotic or biotic responses are specifically associated with increased expression variability. In contrast, the GO enrichment results for low and high NEV groups of singletons (Supplemental Table 5 – 6) and WGDs (Fig. 2C and Supplemental Table 3 – 4) do not exhibit similar patterns observed in SSDs. For example, both low and high NEV groups for WGDs are enriched with central metabolism and biosynthetic processes. These results suggest that the association between elevated NEVs and plastic responses is more pronounced with SSDs than WGDs.

**Figure 2:**
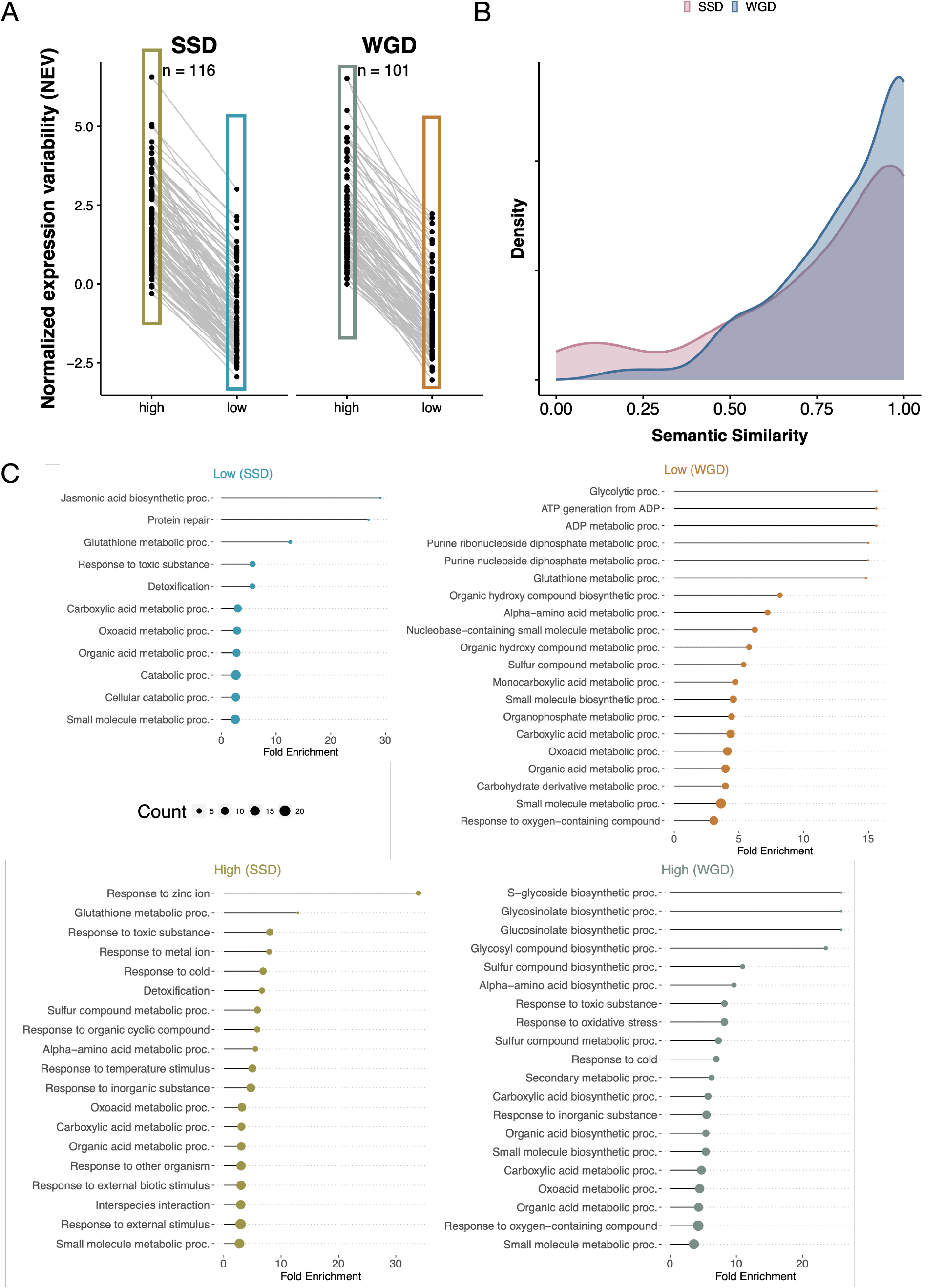
Contrasting GO enrichment results for SSDs and WGDs with NEV diverged gene pairs. A). Top 5% paralog pairs with strongest NEV divergence between two paralogs are shown as single lines for two duplication types (n = 116, SSD; n = 101, WGD). Repeated genes in each group are removed. B). Pairwise semantic similarity analyses are performed for each pair of paralog for SSDs and WGDs. R package GOSemSim (Yu et al. 2010) is used with parameters: measure = “Jiang”, combine = “BMA”, ont = ‘BP’. C). GO enrichment analyses are conducted using ShinyGo http://bioinformatics.sdstate.edu/go/ (Ge et al. 2020), for each group among the four in panel A separately (with corresponding colors). Lists of GO biological functions with FDR of 0.05 are shown. GO terms with less than 10 genes are excluded. Fold enrichment and gene count are shown with lollipop plot with the dot. Enrichment results of singletons with high and low NEVs are shown in Supplemental Table 5 and 6.

We also observed bias in transcription factor retention between two duplication types. The list of TF in *Arabidopsis thaliana* is obtained from PlantTFDB (Jin et al. 2016). We assessed the number of transcription factor (TF) and non-TF paralog pairs in the SSD and WGD pools. (Cases in which a pair of paralogs comprise both a TF and a non-TF gene are rare and thus excluded.) SSDs including proximal, tandem, and transposed duplications have far fewer TF paralog pairs as compared to genes arising from WGD (Fig. S5, p-value = 5.61e-96, chi-squared test), in line with previous results (Maere et al. 2005). Importantly, for tandem and proximal duplications, TF paralog pairs exhibit lower NEVs as compared to non-TF paralogs, and the distribution of NEV between TF and non-TF pairs is more divergent than other duplication types (Fig. S6). This suggests, for small-scale duplications, lower expression variability may be essential for retention among transcription factors.

### Distinct mechanisms leading to elevated expression variability of WGDs and SSDs

Despite comparable elevation in normalized expression variability (NEV) relative to singletons, WGDs and SSDs display distinct functional enrichment patterns among retained genes. We hypothesize that these differences stem from the timing and rate at which high expression variability is acquired. In principle, the *cis*- and *trans*-regulatory environments are likely conserved immediately after WGD events. Consequently, the increased NEV observed in WGD duplicates, relative to singletons, may result from the gradual accumulation of differences over extended evolutionary timescales. In contrast, the elevated NEV in SSDs may emerge during the duplication process itself. Mechanisms such as translocation via transposable elements or reverse transcription can disrupt a paralog’s regulatory machinery, independently of selection or other long-term evolutionary pressures. As a result, the increased NEV observed in SSDs may reflect both the immediate disruptive effects of novel genomic contexts post-duplication and the preferential retention of duplicates with inherently higher NEVs, as those with lower variability may be more prone to early gene loss.

We thus test the gene age bias for SSDs and WGDs. We utilized gene age estimates for 17,732 Arabidopsis genes, inferred based on homologs (a phylostratigraphy approach (Arendsee et al. 2014, Domazet-Loöo et al. 2007)) without paralog information. Many inferred paralogs exhibit differences in their estimated clade of origin (obtained from Arendsee et al. 2014). We assess the gene age bias for using these phylostratigraphy-derived gene age information. The positive trend for SSDs in Fig. 3A and B indicate a bias that the younger gene in a paralog arising from SSD tend to have a higher NEV than the older genes in a paralog. By contrast, no such bias is observed for WGDs in both Fig. 3A and B.

**Figure 3:**
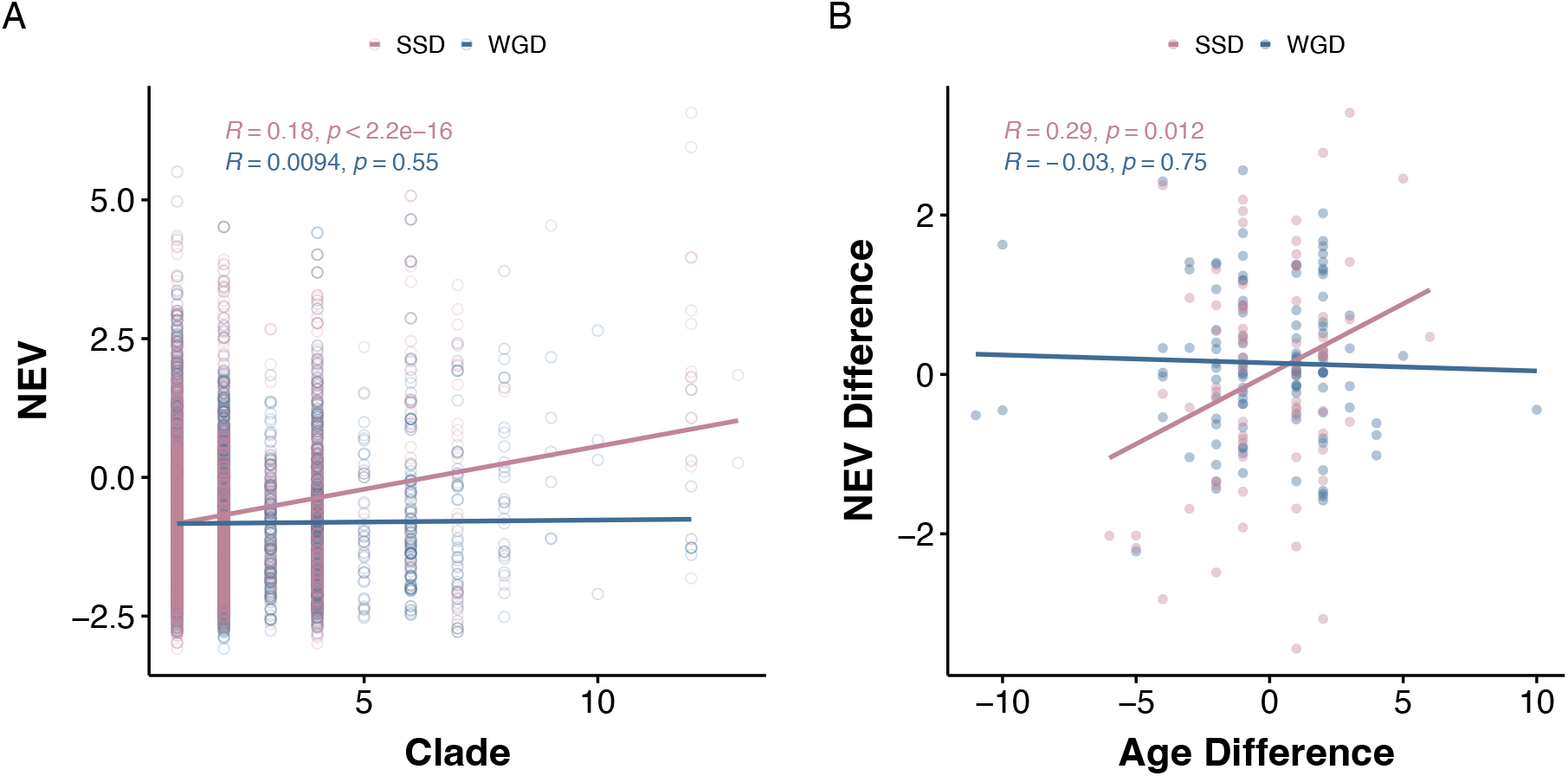
Paralog pairs arising from SSDs and WGDs exhibit distinct age-dependent patterns in NEVs. A). For duplicates arising from SSD, younger genes tend to exhibit higher NEV while no significant patterns observed for WGD duplicates. B). Contrasting patterns between SSD and WGD for gene age and NEV within a paralog pair. In a SSD paralog pair, the younger gene tends to exhibit higher NEV while no significant patterns are observed for WGD paralog pairs. Values in x-axis are the difference of gene clades between paralogs in each pair and the y-axis is the difference of normalized expression variability (NEV) between paralogs in each pair. Paralog pairs with the same inferred clade are excluded (resulting in x = 0). Gene age data were obtained from published data sets (Arendsee et al. 2014), comprising 17,732 Arabidopsis genes stratified into 15 clades, numbered from the oldest (stratum 1) to the most recent (stratum 15).

We speculate that the discrepancy of gene age between paralogs derived from phylostratigraphy approach, to some extent, indicates the time of origin for a paralog copy. The relative timing of origin can further reflect which paralog is the parental copy and which one is the daughter copy. To validate this hypothesis, we used an existing data of transposed duplication paralogs (a subset of SSDs), which specifies which paralog is the parental/daughter copy via a synteny approach — multiple colinearity scans (MCScanX) (Wang et al. 2012a, 2013). The results indeed show that, daughter loci derived from synteny analyses tend to be the younger copy (paired Wilcoxon test p-value = 1.035e-06; sample size n = 1373).

In addition, our analyses revealed consistently distinct temporal patterns for NEV divergence between SSDs and WGDs, when estimating duplication times with two independent methods (Fig. S10, Fig. S11, and Fig. S12). Though, within either duplication mode, the observed patterns were method-dependent and require further investigations.

Our findings support distinct models underlying elevated NEV in SSDs and WGDs. WGDs show no age bias, whereas SSDs exhibit a bias toward higher NEV for the younger genes (Fig. 3B), driving the overall positive relationship between gene age and NEVs for all genes (Fig. S9). Our follow-up analyses demonstrate that the gene age can be used to identify parent/daughter in a paralog. Taken together, this suggests that for SSDs, the daughter copy in a paralog tends to have higher NEVs than the parental one.

## Discussion

Although the direction of causality between expression variability and gene retention cannot be determined conclusively, our analyses suggest a plausible model to explain the elevated expression variability observed for SSDs and WGDs. Translocation of a paralog to a novel genomic setting is likely to lead to regulatory changes, resulting in expression variability differences between the new paralogs. SSDs that arise from unequal crossing over during meiosis may not faithfully duplicate the complete *cis*-regulatory sequences, leading to mis-regulation as compared to the ancestral single copy gene (Rogers et al. 2017). SSDs that arise from translocation, e.g. via transposable elements or reverse transcription, may likewise find themselves in novel genomic contexts where local features such as chromatin structure (e.g., topological association domain), methylation, or *cis*-elements introduce novel expression phenotypes (Wang et al. 2012b). The higher NEV for paralogs can be due to two factors that are not mutually exclusive: First, the immediate regulatory changes tend to increase NEVs. Second, there is no bias in NEVs with novel genomic context post-duplication. However, those new duplicates with increased NEVs may have higher retention probability.

In line with this model, we observe a bias towards higher NEV for daughter genes in SSDs (Fig. 3B). Indeed, Qiao et al. 2019 synthesized duplication data across plant lineages and reported that proximal and tandem duplicates exhibited relatively high *K*_*a*_/*K*_*s*_ ratios but small *K*_*s*_ values, possibly a signature of more rapid functional divergence as compared to other gene classes and supporting the hypothesis that positive selection plays an important role in the early stages following small-scale duplications (SSDs). In contrast to SSDs, we observe opposite patterns for WGDs in Fig. 3. We speculate that the overall elevated expression variability for WGD could potentially be due to relaxed selection due to extra copies or positive selection (Zhang et al. 2009) over prolonged evolutionary time.

As such, stress-responsive genes are more likely to be retained via SSD than via WGD. Conversely, the retention of transcription factors follows a different pattern, for which dramatic regulatory and expression variability changes are likely deleterious as these can be central and highly ‘connected’ in gene regulatory networks and therefore subjected to selective pressure to minimize transcriptional noise (PuzoviÊ et al. 2023). Therefore, transcription factors are less likely to be retained following SSDs than WGDs. Our results propose a plausible explanation for reciprocal retention between tandem duplication and WGDs, potentially complementary to dosage balance theory and other existing models (Freeling 2009, Tasdighian et al. 2017).

The concept of inter-individual variability we study here is distinct from the expression noise in single-cell organisms, which is cell-to-cell heterogeneity in gene expression. The cell-to-cell heterogeneity in gene expression is due to the stochastic nature of the molecular interactions involved in (post)transcription regulation (Elowitz et al. 2002, Raser and O’shea 2005), including variation in the internal states such as the abundance of key cellular components (Pedraza and van Oudenaarden 2005, Das Neves et al. 2010), and external random variation such as micro-environment variation among genetically-identical cells (Battich et al. 2015). Besides these sources of variation, the inter-individual expression variability can arise from micro-environmental variation among individuals (during development) and stochastic variation in cell-type compositions among individuals (bulk RNA-seq experiments measure total gene expression from heterogeneous tissues). Micro-environmental variation, which leads to cell-to-cell heterogeneity and inter-individual variability, is distinct from macro-environmental variation, which regulates stress responses and phenotypic plasticity (Hall et al. 2007, Siegal and Leu 2014, Paaby and Testa 2021, Cortijo et al. 2019). In experimental settings, macro-environmental variation refers to systematic environmental differences deliberately introduced by the experimenter, whereas micro-environmental variation encompasses mild, non-directional random fluctuations (Masel and Siegal 2009).

Low genetic variation is often interpreted as an evolutionary constraint. In many cases, the within-line phenotypic variability and noise are associated with genetic variation within population as well as environmental plasticity - the so-called congruence hypothesis (Wagner et al. 1997, Gibson and Wagner 2000, Hansen 2006). One explanation for congruence is that different causes of perturbations may be aligned because the responsive dynamics reflect the intrinsic tendencies of developmental systems to produce certain forms rather than others (Alberch 1989, 1982, Nuño de la Rosa and Müller 2024, Landry et al. 2007). Our study demonstrates a case where such variation, apparently arising from stochastic factors, may be used as an index of evolvability (Draghi and Ogbunugafor 2023, Rocabert et al. 2020). Our study emphasizes the role of expression variability in shaping environmental plasticity and expression divergence, pointing to the need to assess phenotypic noise in evolutionary studies (Cai et al. 2024).

## Acknowledgements

The authors thank three anonymous reviewers, Matt Rockman, Xiang Li, Robin Hopkins, Chuliang Song and Des Marais Lab members for helpful discussions. This work was supported by NSF IOS 2239070 and a grant from the Robert and Ardis James Foundation to DLD.

